# Kinematics of individual muscle units in natural contractions measured *in vivo* using ultrafast ultrasound

**DOI:** 10.1101/2022.05.26.493608

**Authors:** Emma Lubel, Bruno Grandi Sgambato, Deren Y. Barsakcioglu, Jaime Ibáñez, Meng-Xing Tang, Dario Farina

## Abstract

**Objective:** The study of human neuromechanical control at the motor unit (MU) level has predominantly focussed on electrical activity and force generation, whilst the link between these, the muscle deformation, has not been widely studied. An important example of this is excitation-contraction coupling (E-C coupling) – the process by which electrical excitation is converted into contraction in the muscle fibres. Despite this being a clear marker for progression of certain diseases, it cannot be measured *in vivo* in natural contractions. To address this, we analyse the kinematics of muscle units in natural contractions.

**Approach:** We combine high density surface electromyography (HDsEMG) and ultrafast ultrasound (US) recordings of a mildly contracted muscle (tibialis anterior) to measure the deformation of the muscular tissue caused by individual MU twitches (decomposed from the HDsEMG). With a novel analysis on the US images we identified, with high spatio-temporal precision, the velocity maps associated with single muscle unit movements. From the individual MU profiles obtained from the velocity maps the region of movement, the duration of the mechanical twitch, the total and active contraction times, and the activation time (equivalent to E-C coupling) were computed.

**Main results:** The E-C coupling was 3.8 ± 3.0 ms (n = 390), providing the first measurement of this value in for single MUs in non-stimulated contractions. Furthermore, the experimental measures provided the first evidence of single muscle unit twisting during voluntary contractions and showed the presence of MUs with territories with multiple distinct split regions across the muscle region.

**Significance:** We show that the combined use of HDsEMG and ultrafast US can allow for the study of kinematics of individual MU twitches, including measurement of the excitation-contraction coupling time under natural neural control conditions. These measurements and characterisations open new avenues for study of neuromechanics in healthy and pathological conditions.

## Background

Motor units (MUs) translate synaptic input into muscle motion: the action potentials (Aps) discharged by the spinal motor neurons (MNs) in response to synaptic input evoke transient mechanical contractions, called twitches, of the fibres of the muscle unit. Ultimately, global motion is a result of the summation of these muscle unit twitches. The neuromechanics of movement can therefore be studied at the MU level, where the ultimate neural code of movement is translated into mechanical output.

Human neuromechanical control is well explored from both an electrical perspective, providing little spatial detail, and in terms of muscle contraction, mainly treating the muscle as a global force producer. However, how the microstructures in the muscle combine to do so is less clear – the complexity and degree of linearity of the system are unknown. A representative example of a mechanism relevant for understanding the neuromechanics of muscle control is excitation-contraction coupling (E-C coupling), which is one of the components of the electromechanical delay (EMD). E-C coupling refers to the communication between the electrical activity in the skeletal muscle fibre membrane and the release of calcium, eliciting the contraction (Calderón, Bolaños and Caputo, 2014). The ability to measure E-C coupling *in vivo* in voluntary contractions for individual MUs would provide insights into neural control as the process of modulating MU activities, and would also enable detection and monitoring of pathological conditions in which the E-C coupling is affected, such as myotonic dystrophy 1 (Esposito *et al*., 2016) and lower back pain (Vasseljen *et al*., 2006). Coupling time has been measured in electrically stimulated contractions (Nordez *et al*., 2009; Esposito, Limonta and Cè, 2011; Hug *et al*., 2011) or in voluntary contractions at the global muscle level with problematic reliance on bipolar EMG (Hug, Lacourpaille and Nordez, 2011; Begovic *et al*., 2014). A method to simultaneously study the electrical and mechanical behaviour of active MUs is required.

Generally, however, methods for MU characterisation can be divided into two broad categories – those which consider the electrical activity, and those which consider the mechanical output. The electrical behaviour of MNs is usually analysed with electromyography (EMG), either intramuscular (iEMG) or surface (sEMG). Both iEMG and sEMG allow the indirect identification of MN discharges by the analysis of the corresponding muscle unit APs. On the other hand, the mechanical response of entire muscles can be studied using ultrasound (US) (Botter *et al*., 2013; Dieterich *et al*., 2017; Tweedell, Tenan and Haynes, 2019). US has also been used for studying the mechanical response of individual MUs in electrically stimulated contractions (Deffieux *et al*., 2008; Waasdorp *et al*., 2019, 2021). Furthermore, some studies have suggested that US can even be used at the muscle unit level (Rohlén *et al*., 2020; Rohlén, Stålberg and Grönlund, 2020; Carbonaro *et al*., 2022). Here, we utilise US alongside a new signal processing pipeline, with minimal processing and assumptions, to extract detailed new characteristics of the kinematics of single muscle units.

In this work we used ultrafast US to extract velocity-time profiles for twitches of MUs. From these we characterised the isolated movement of muscle units. We describe and validate the proposed methodology, and we provide results on new physiological measurements not previously possible, including MU-specific E-C coupling time, allowing deeper insight into neuromechanics. We also present the first evidence of single MU twisting during muscle twitches.

## Methods

### Ethical approval

All procedures and experiments were approved by the Imperial College Research Ethics Committee (ICREC reference: 20IC6422) in accordance with the declaration of Helsinki. As the datasets acquired are large, the data was not registered in a public database but may be made available, as appropriate and reasonable, upon request.

### Experimental Setup and Equipment

The method described is based on synchronised acquisition of ultrafast US, high-density surface electromyography (HDsEMG), and force data during repeated sustained contractions performed with feedback on MU activity provided by real-time HDsEMG decomposition. Figure 1 summarises how APs generate contractions and shows the information captured by each modality.

**Figure 1.**
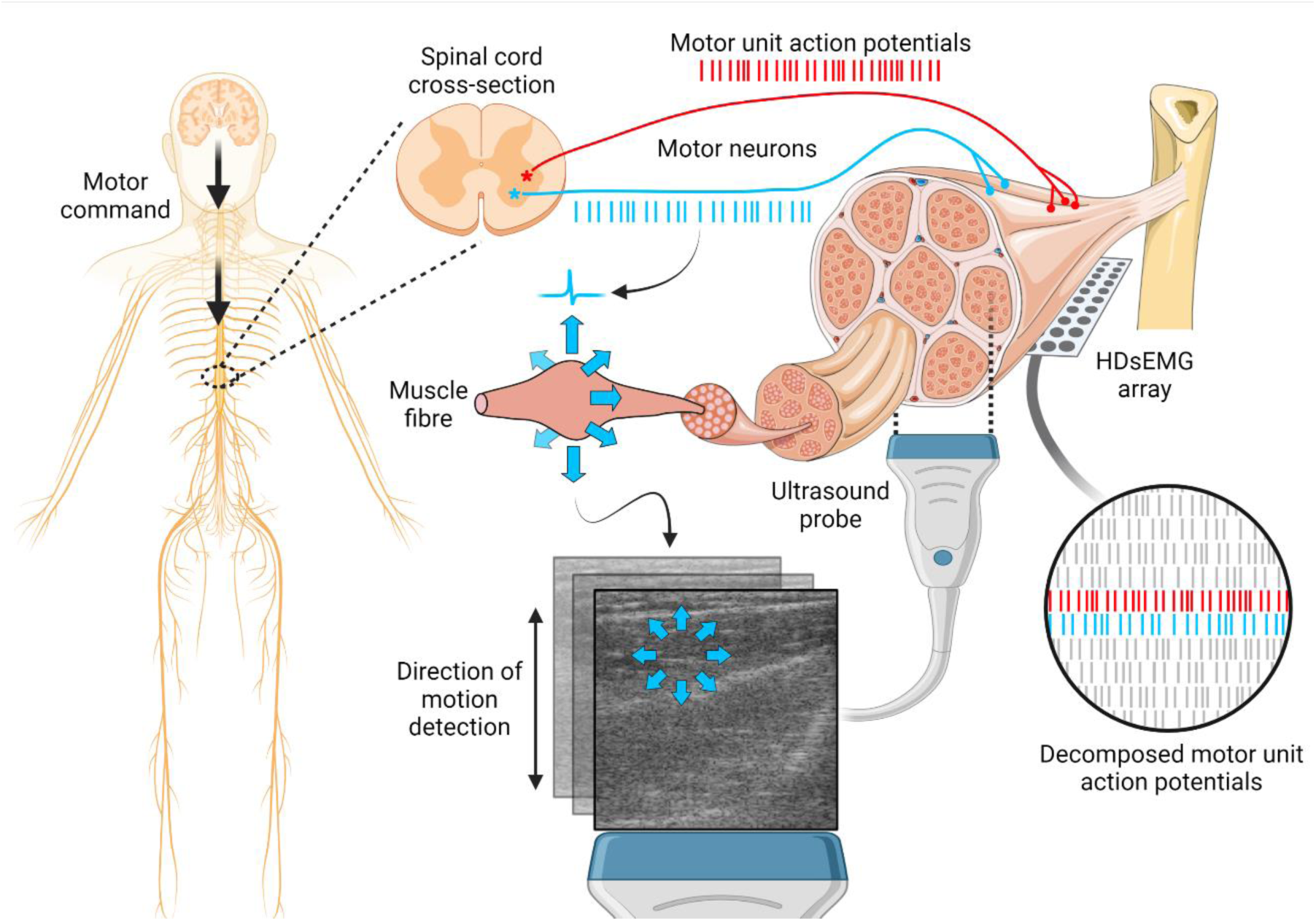
Diagram of the experimental set-up and the underlying human neuromechanical control mechanisms. Motor command is passed from the brain via the spinal cord to the motor neurons (MNs). Each MN innervates a group of muscle fibres. The ensemble of action potential (AP) trains from the pool of MNs innervating a muscle are often referred to as the neural drive to the muscle. Together, the innervated muscle fibres (muscle unit) and the motor neuron constitute a motor unit (MU). APs travel along the MN axon and generate corresponding APs along the muscle fibre membranes. These cause a contraction (shortening and thickening) of the fibres. The ultrasound (US) imaging plane is perpendicular to the direction of the fibres, so a radial expansion is seen in the imaging plane. The velocity mapping algorithm detects axial motion only, so only the components of the expansion perpendicular to the probe are detected. High-density surface EMG (HDsEMG) arrays (for clarity, only one is shown in this diagram) are placed over the surface of the skin, and an online decomposition algorithm is used to estimate the motor unit action potential (MUAP) discharge times.

#### Ultrafast US

The US data were acquired with the Vantage Research Ultrasound Platform (Verasonics Vantage 256, Kirkland, WA, USA), equipped with a L11-4v transducer with 128 elements and centre frequency of 7.24 MHz. Single angle plane wave isonification was used to enable a frame rate of 1000 frames per second with acquisitions of 30 seconds, resulting in 30,000 frames of US data per recording. The data was beamformed using delay and sum beamforming, resulting in images of 357 pixels axially and 128 laterally, corresponding to pixel dimensions of 0.1 and 0.3 mm. The frame rate and recording duration were chosen as per memory limitations. The chosen values proved to be a good compromise between temporal resolution and duration for accurate processing. The acquisition and control were performed by custom codes written in MATLAB (Mathworks, Massachusetts, USA).

#### High-Density Surface Electromyography

The HDsEMG signals were recorded by two electrode arrays of 64 channels (5 columns and 13 rows; gold coated; 8 mm interelectrode distance; OT Bioelettronica, Torino, Italy). The signals were recorded in monopolar derivation, amplified, sampled at 2048 Hz, A/D converted to 16 bits with gain 150, and digitally bandpass filtered (10-500 Hz), using an EMG pre-amp and a Quattrocento Amplifier (OT Bioelettronica, Torino, Italy).

#### Force

An ankle dynamometer was used to measure the dorsiflexion force. The leg and foot were constrained in the dynamometer using foam padding and secured using straps to provide stability and to eliminate other motions, keeping the foot at an angle of 90 degrees to the leg. The force data was fed through a Forza force amplifier (OT Bioelettronica, Torino, Italy) directly into the auxiliary port of the Quattrocento amplifier.

#### Synchronisation

A 1 µs active low output was fed from the Verasonics US system into an Arduino UNO that in turn outputs a 490 µs pulse, elongated to allow for detection at the 2048 Hz sampling frequency of the EMG amplifier. The trigger was recorded at the start and end of each recording period and used to align the data in post-processing. To test the accuracy of synchronisation and alignment of the US series and the HDsEMG signal, the HDsEMG grid was sharply perturbed such that a peak was seen in the electrical (as artefact) and velocity signal obtained from the US series. These peaks were seen to align within the expected error of ± 0.5 ms resulting from the US frame rate, confirming the validity of the synchronisation method.

#### Participants

A total of 12 healthy participants (mean age, 26.25 ± 2.8, 4 female), were recruited for this study. Before the experiment, the volunteers were fully briefed on the experiment, provided with a participant information form, and asked to sign an informed consent form.

#### Setup

The experiment was conducted on the tibialis anterior (TA) muscle. The motivation for using the TA in this study was threefold: it is a large, long muscle allowing space for all the sensors used; it is superficial, resulting in high-quality HDsEMG decomposition (Del Vecchio *et al*., 2020); and its fibres are long and have a relatively low pennation angle (Sopher *et al*., 2017) (see figure 2). The latter property allowed for mounting the ultrasound probe and the HDsEMG arrays in separate areas whilst ensuring they still overlay the same fibres, and ensuring more uniform velocity fields within the imaging plane. In all participants, the experiment was performed on the (dominant) right leg.

**Figure 2.**
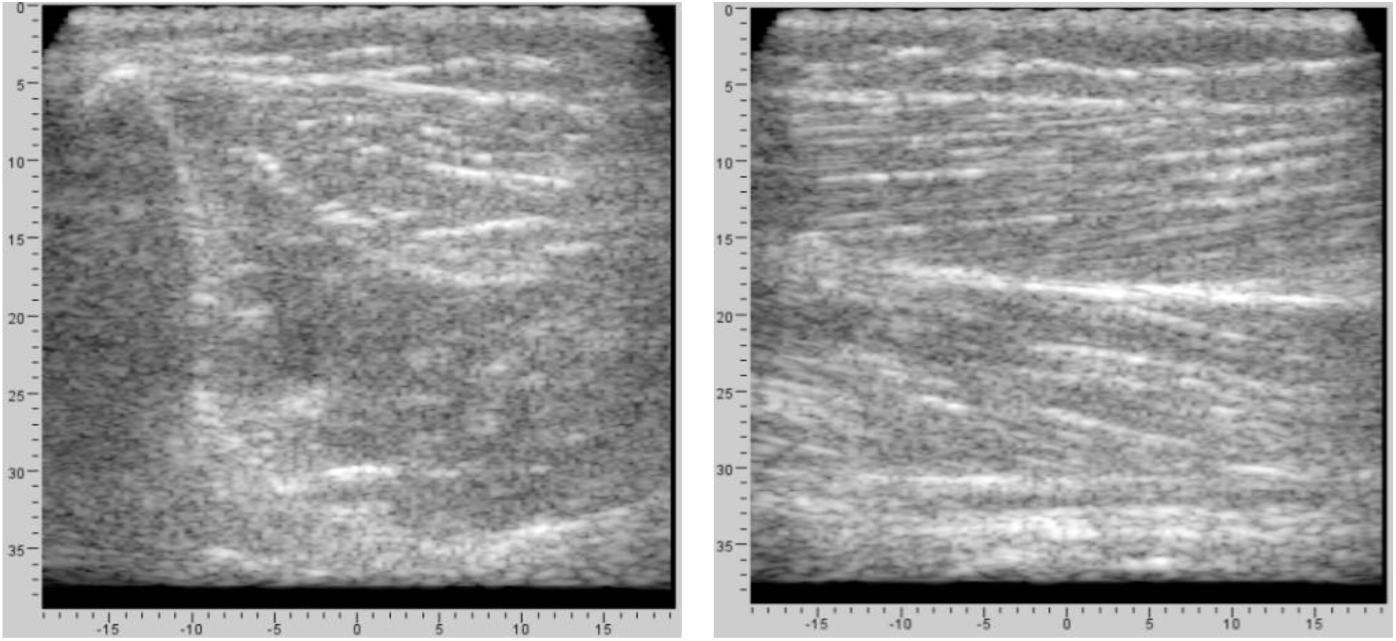
Coronal (left) and parasagittal (right) ultrasound images of the tibialis anterior (TA) muscle at the point of recording used in the experiment. From the right hand image it can be seen that the low pennation angle of muscle fibres results in fibres almost parallel to the skin.

Before beginning the experiment, the skin over the TA was shaved, lightly treated with a chemical abrasive, and cleansed using an alcohol spray. The muscle belly was then identified by palpation whilst the participant was guided to contract and relax the TA. A HDsEMG electrode array was placed over the proximal part of the muscle, parallel and lateral to the tibia. The grid was positioned such that all electrodes were on top of the TA belly. A 1.5 cm gap was left distally with respect to the first array and a second array was placed over the distal part of the muscle, also parallel and lateral to the tibia. In some participants, the distal electrodes of the second array fell outside the TA belly, which resulted in small signals and did not impact the results. The electrodes were placed along the estimated direction of the muscle fibres and secured using Tegaderm Film Dressings and medical self-adhesive bandages.

Next, the US probe was placed in the gap between the EMG electrodes, with the imaging plane perpendicular to the length of the muscle fibres. A water-based US gel was used to improve the coupling between the probe and the skin. The US probe was held by a custom-built probe holder and secured in position using straps. The participant sat in a chair in a comfortable position with the leg strapped into the ankle dynamometer. The US system was turned on to perform real-time imaging and small adjustments were made to ensure the TA muscle belly was covered. Finally, a computer screen was adjusted to a comfortable distance and position to provide visual feedback during the experiment. The described configuration of separate locations for the EMG grids and US probe was preferred over the possibility of using grids transparent to US directly under the US probe (Botter *et al*., 2013) since transparent grids have not been tested for single muscle unit motions (see also Discussion).

### Experiments

Real-time online decomposition of the HDsEMG signals into MU discharge times was performed using the methodology of Barsakcioglu et al. (Barsakcioglu *et al*., 2021). The identified spike trains were provided as visual feedback to the participant throughout the experiment. This approach ensured continuous activation of the target MUs, and allowed the participant to precisely control the number of recruited MUs.

The online decomposition was trained on a contraction at 10% maximum voluntary contraction (MVC), maintained for 41 s. Once the online decomposition model was trained, the subject was asked to slowly increase the dorsiflexion force whilst the experimenter inspected the decomposition to determine whether the model was satisfactorily trained, and the noise level was acceptable. Model training was repeated when deemed necessary (this was never done more than once during the experiments).

With the decomposition trained, the visual feedback displayed the MU discharge times in real-time. The subject was asked to slowly increase the dorsiflexion force, thereby slowly increasing the number of recruited MUs displayed on the screen. The order of recruitment of the identified MUs was noted. The subject was then trained to recruit individual MUs and to incrementally increase the number of MUs in a controlled manner. Providing visual feedback was a vital part of the training as fine recruitment of low threshold MUs is a complex task.

Once the participant was comfortable with this control method, the recordings commenced. The subject was instructed to recruit a single MU and to keep it active for 30 s whilst the US and HDsEMG recordings were taken. The subject then rested for 60 s before repeating this process for the same MU two further times. Each 30 s paired recording of HDsEMG and ultrafast US was considered a single trial. The three trials were then repeated for up to six different MUs. If fewer than 6 MUs were identified, the process was only performed up to the maximum number of identified MUs.

### Data Analysis and Processing

#### Datasets

For one subject the synchronisation signal failed, and for another the HDsEMG decomposition failed, resulting in valid data from 10 subjects. For an example participant, 8 MUs were detected by HDsEMG and 18 paired recordings of HDsEMG and US were taken. Of these 8 MUs, 6 were in the correct plane to be seen by the ultrasound probe, resulting in 12 usable paired recordings. In each of these 12 recordings multiple MUs were active, and the number of active MUs varied. As such, from these 12 recordings, 33 MU-STAs were computed. Overall across 12 participants 51 MUs were identified in both US and HDsEMG and 390 MU-STA curves were obtained.

#### Data Processing

From the online decomposition, the discharge times of the identified MUs were available. However, each discharge was identified with some variability with respect to the AP waveform. Since in the current application it was important to define the precise onset of the MU discharge, we estimated the required temporal shift so all APs would be aligned in time. The shift was identified by means of a STA of the HDsEMG signals over each channel, producing the average AP shape for each MU. A double differential AP shape was calculated and the channel with highest signal-to-noise ratio was selected, and the onset of the action potential was defined as the first time sample with amplitude higher than 5 standard deviations of the baseline signal (Ibáñez *et al*., 2021). The time difference between the onset and the centre of the STA window was the shift by which all AP spikes of the MU were shifted, such that the corrected AP discharge time reflected the onset of electrical potential at the neuromuscular junction in a consistent manner for all discharges.

For the ultrafast US data, each trial comprised 30,000 frames (357 × 128 pixels each) that were used to generate a series of velocity maps using 2D autocorrelation velocity tracking (Loupas, Gill and Powers, 1995) with a sliding window of 2 frames, corresponding to 2 ms. This resulted in 29998 frames (357 × 128 pixels) with bipolar amplitude representing axial velocity (negative pointing away from the probe). The recordings were then further low-pass filtered (0.5 MHz) along the axial direction and band-pass filtered (5 – 100 Hz) along the temporal direction, removing noise from outside the expected temporal range of the response.

Next, a STA of the US velocity maps was performed using the discharge times identified from the HDsEMG decomposition. For each identified MU, a ± 50 ms window (100 frames in total) of velocity data around each discharge was selected (figure 3). All windows were averaged, resulting in 100 frames of STA US maps (US-STA maps) for each MU (figure 4(a)).

**Figure 3.**
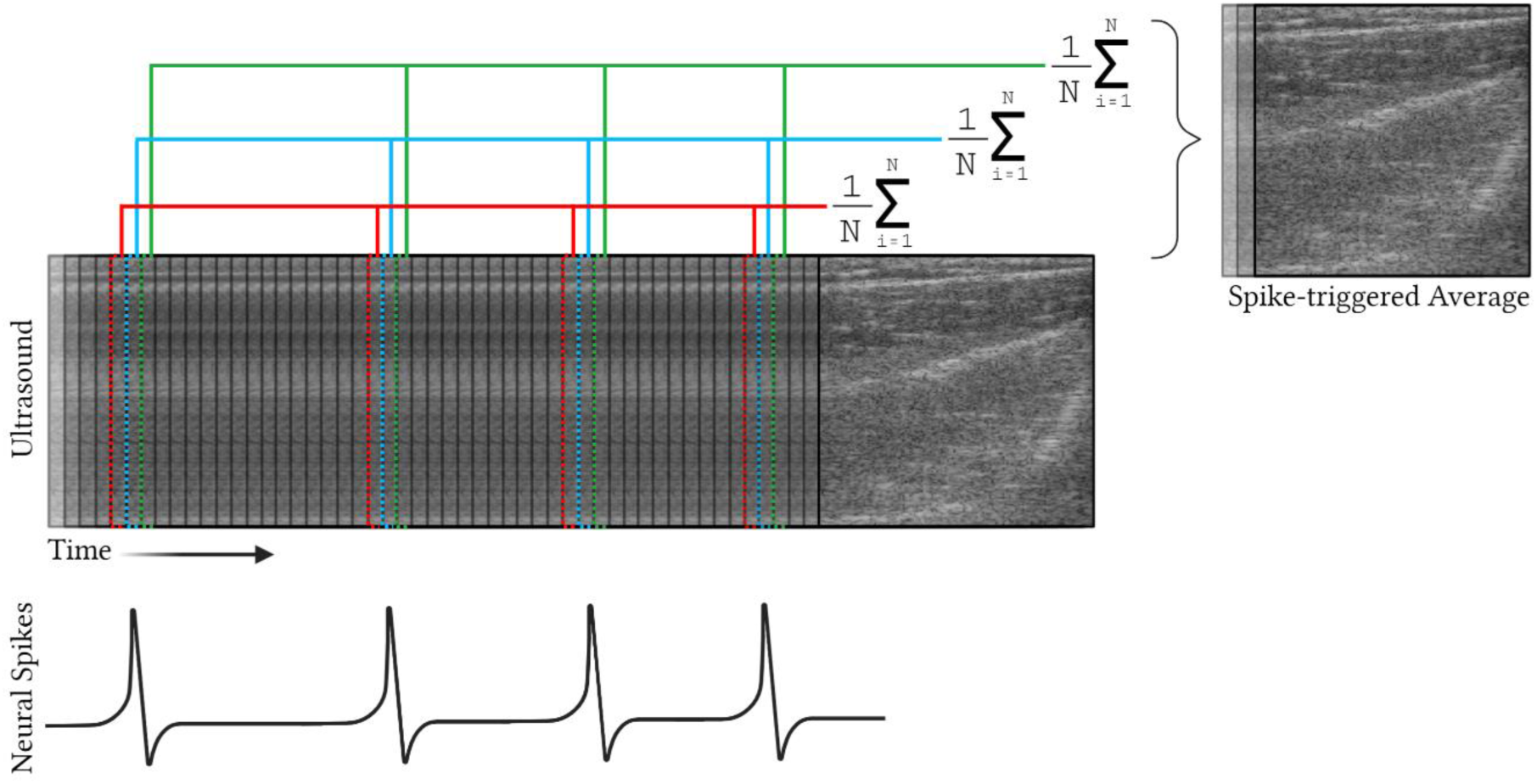
Diagram of the spike-triggered average (STA) method employed. For each motor unit (MU) and identified discharge, a window around the discharge timing is selected. All selected windows are summed and averaged, resulting in a single STA window for each MU. Note that this process is performed on the velocity fields obtained from the ultrasound image series.

**Figure 4.**
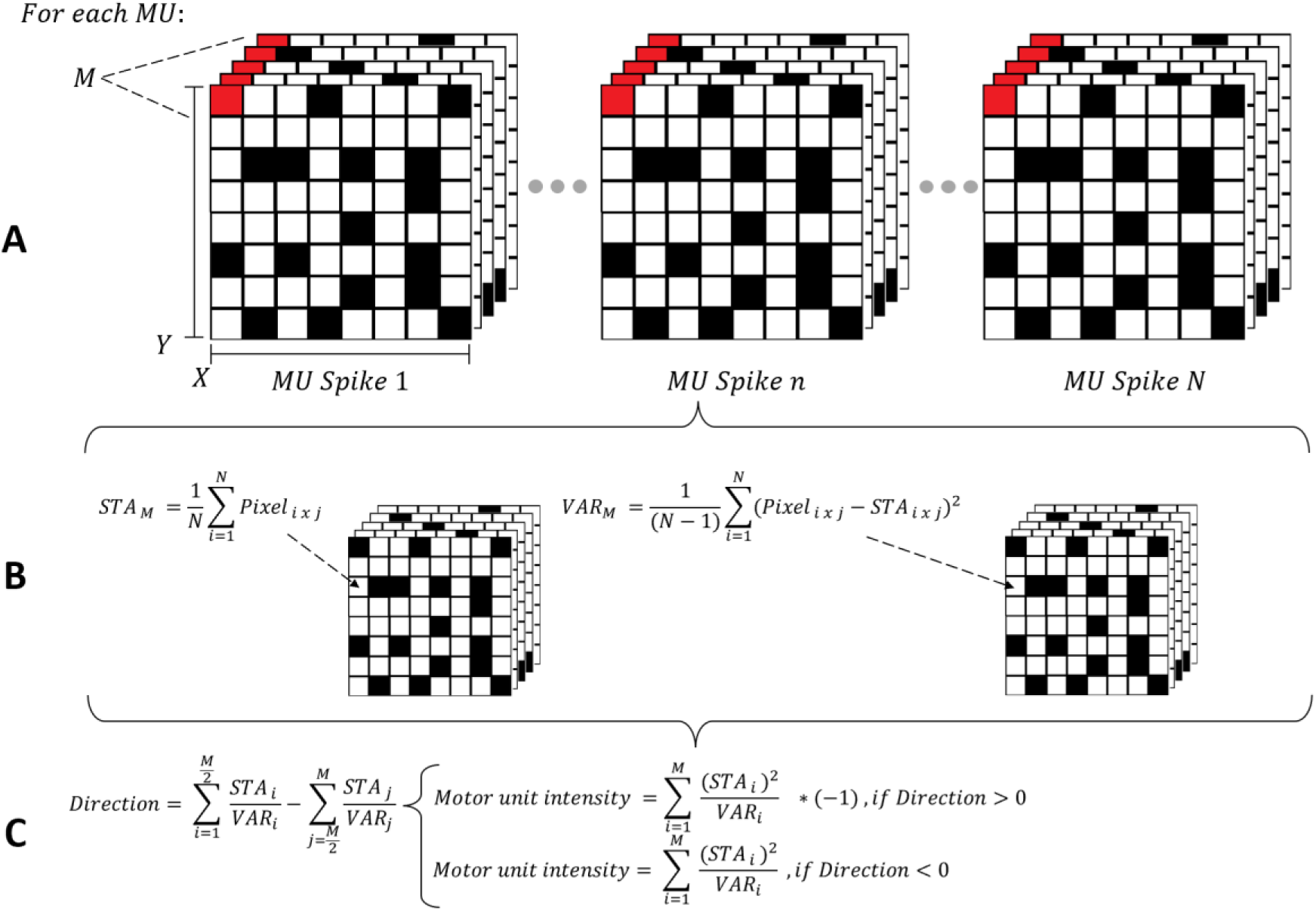
Process to highlight motor unit (MU) related motion. For each MU a window of 50 frames before and 50 frames after each action potential (AP) time is selected (shown in panel A, where M = 100 frames, N = total number of APs in that recording, X = width of the image, and Y = depth of the image) and a spike-triggered average (STA) technique is employed to obtain the STA and the variance of the STA, as shown in panel B. The STA is divided by its variance and summed over time to obtain a single map of intensities, where high intensity represents areas of high motion and low variability in the STA. The values are also multiplied by a direction factor (as shown in panel C) which separates segments of the STA map moving in opposite directions in response to the AP.

Supplementary Video 1 in the Supporting Information shows an example of the 100 frame US-STA i.e. the average muscular movement caused by an individual MU discharge. In this representative video, a large part of the movement caused by the target MU is hidden by the motion of non-muscular structures. These high noise areas need to be removed for an accurate analysis of the muscle unit movement. For this purpose, for each US-STA window, the variance across all windows was calculated (figure 4(b)). To determine the direction of motion of a pixel as induced by the MU activity (either positive – towards the probe – or negative – away from the probe), we performed a subtraction of the sum of the second half of the STA from the first (figure 4(c)). For each pixel the sum over the squared STA divided by the variance (with the convention of a positive sign for upwards and negative sign for downwards motions) was calculated, highlighting regions of intense (high amplitude) and consistent (low variance) movement, whilst penalising regions of non-consistent movement (e.g., tibia movement). Lastly, the map was downsampled axially by a factor of 3 (119 × 128 pixels wide; 0.3 × 0.3 mm per pixel) and the regions of no interest (the rows after the maximum sEMG range) were removed. Each of the final maps, which we will term MU activity maps (MUAMs) (55 × 128 pixels wide), represents the muscular movement caused by an individual MU discharge during a voluntary isometric contraction.

#### Muscle Unit Area of Motion Identification

The MUAMs show, for each pixel, the effect caused by the discharge of a specific MU, with high values (positive or negative) representing regions of large motion that are highly synchronised with the MU discharge, whilst low values represent regions with either small or highly desynchronised motion. In some situations, a MUAM may show separate regions of positive and negative velocity associated with a single muscle unit motion. In these cases, the two regions were treated separately but assigned to be part of the same MU. To properly define which pixels were associated to the movement of a muscle unit, a thresholding approach was used. For each MUAM, pixels with absolute value greater than 65% (chosen empirically) of the maximum amplitude were considered as part of the region of movement due to the MU activity. We will refer to the pixels constituting this region as the MU motion domain. In contrast to the MU territory, defined as the cross-sectional area containing the muscle fibres of a MU, the MU motion domain is therefore defined as the region of muscle tissue that shows movement with amplitude and synchronicity over a predefined threshold when a MU discharge occurs. Whilst the MU territory refers to the MU anatomy, the MU motion domain refers to the mechanics of the muscle unit.

#### Motor Unit Mechanical Twitch Profile

The MUAMs were used with the US-STAs to generate a STA of the mechanical twitch of the MU (MU-STA curves). These were calculated by averaging the US-STA pixels curves inside the MU motion domain. One MU-STA curve twitch profile was generated for each MU (two if the MU caused both a positive and negative movement in separate regions) in each trial. In this work, twitches refer to the movement of the fibres in terms of a displacement velocity.

Each extracted MU-STA curve is a unique twitch profile that was characterised the parameters shown in figure 5(a). Limited work in this area has resulted in unclear and inconsistent naming of parameters and, as such, our nomenclature may not be in agreement with other works. One extracted feature was the duration of the mechanical twitch – the time over which the tissue drastically changes its velocity profile. This corresponds to the time interval over which the tissue accelerates, thus the boundaries of this interval are inflection points. In some cases (figure 5(b)) the onset of the mechanical twitch can be easily identified by the inflection point of the velocity, however in other cases (figure 5(c)) the twitch occurs during a later stage of relaxation of the previous twitch, so although there is a sharp velocity change, the velocity inflection point cannot be used to define twitch onset. As such, the peak in the second derivative was used to mark the onset of the mechanical twitch.

**Figure 5.**
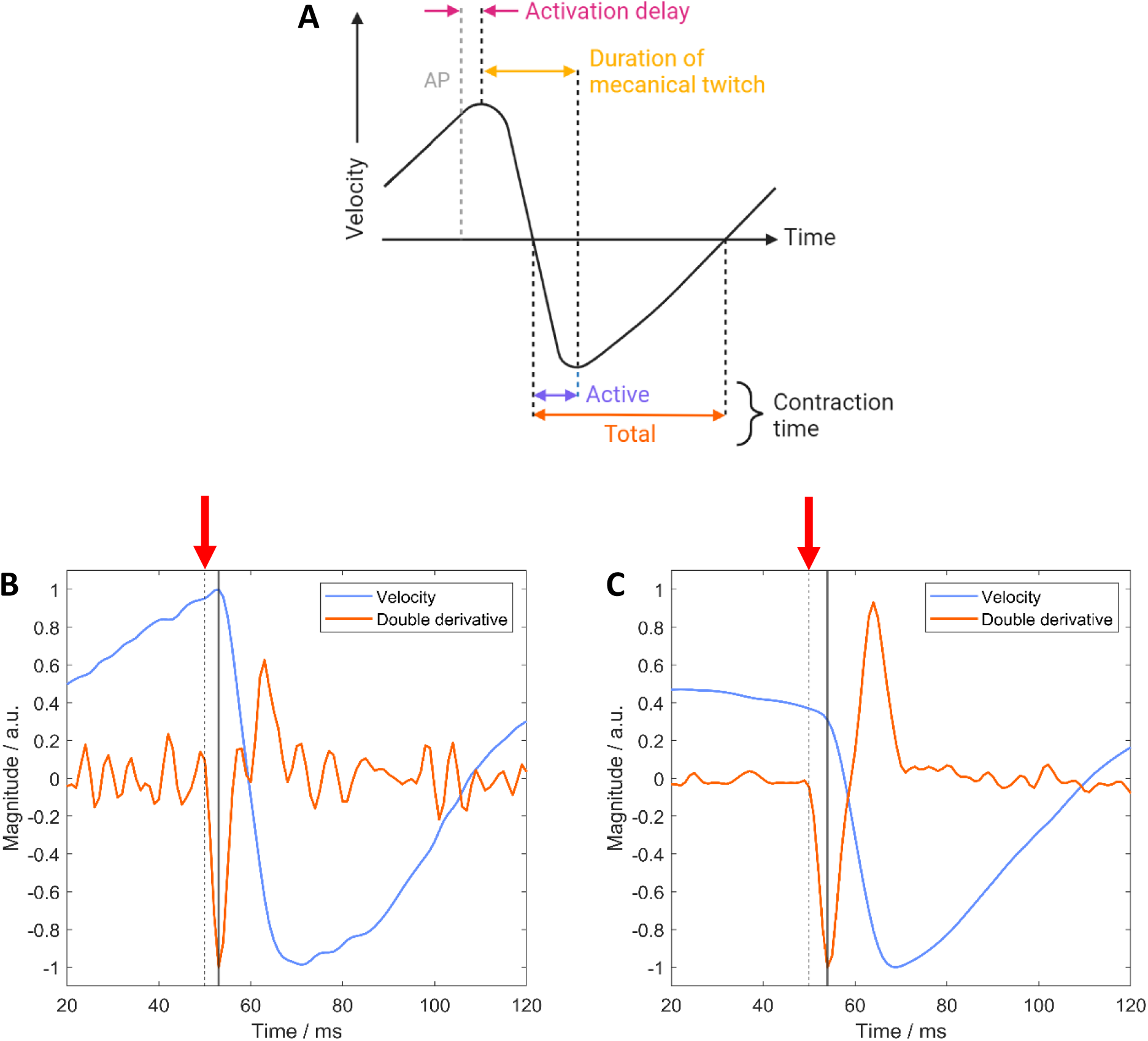
Measurements extracted from the motor unit spike-triggered average (MU-STA) curves. A) The dotted AP line is the time of the onset of the electrical action potential (AP) measured using high-density surface EMG. Note that before the effects of the next twitch begin, the fibre is still relaxing from the previous twitch, thus the velocity is non-zero and changing prior to the AP. Following the AP, the velocity reverses due to the twitch, and the time between the AP and the turning point is the activation delay. The fibre is then accelerating due to the twitch between the first and second inflection points – we refer to this as the duration of the mechanical twitch. Between the two zero crossings, the fibre is in a contracted state – thus we break this region down into active contraction time, where the fibre is being actively contracted, and total contraction time, which also accounts for the relaxation back to the neutral position. B) An example velocity profile for a motor unit (MU) where the interspike interval is low, so an inflection point is seen at the onset of the mechanical twitch. C) An example velocity profile for a MU where the interspike interval is high, so the previous mechanical twitch is in a later stage of relaxation resulting in no turning point at the onset of the twitch. To enable twitch onset detection in both cases, the peak of the second derivative is used (solid black line). In each case, the dotted black line is the firing time from the EMG, highlighted by a red arrow above the plots.

Another feature of interest was the contraction time, defined as active or total. The active contraction time was defined as the interval between velocity zero crossing and the inflection point. The total contraction time starts at the same point, however it ends at the second zero crossing before the next spike. Although both features represent the time where the tissue is in the “contracting state”, the active contraction time accounts for the time over which the twitch is actively accelerating the tissue, whereas the total contraction time includes the full duration of the “contracting state” – the total contraction time includes both the time when the fibres are actively generating the contraction, as well as the time when the acceleration is due to the elastic recovery forces.

Finally, we also calculated the activation delay, defined here as the time between the onset of the electrical AP and the onset of the mechanical twitch. Currently, the electromechanical delay (EMD) is measured as the delay between the onset of the detected electrical activity and the onset of force production, and often its measure has significant inaccuracies due to the use of bipolar EMG at a distance from the motor point (Cavanagh and Komi, 1979). As such, in literature, the EMD includes the time lag due to the synaptic transmission, the excitation-contraction (E-C) coupling, the muscle force transmission on the series elastic components, the aponeurosis, and force transmission on the tendon. Our technique allows for the measurement of the delay between the onset of voluntary motor unit action potentials (MUAPs) and the first measurable acceleration on the fascicles of single muscle units. Therefore, this measure includes the time lag caused by the E-C coupling and force transmission on the series elastic components (between the innervation point and the US measurement plane). However, as the force transmission travels at ∼30 m/s (Morimoto and Takemori, 2007; Nordez *et al*., 2009) and the end-plates for the detected MUs were within 4-cm distance to the US plane, the resulting delays were less than 1 ms. Thus, the measured activation delay mainly represented the E-C coupling, making our measurements the first to report individual MU E-C coupling delays.

#### Processed Dataset

Following the process described above, all 179 paired recordings were processed. However, 31 recordings were discarded as the HDsEMG decomposed spike trains were poorly identified. A total of 68 MUs were identified across all participants in the HDsEMG recordings, 51 of which (75%) could also be detected in the US recordings. This discrepancy is due to out of plane MU activity, a problem which is further explored in the Discussion. As such, 390 MU motion domains and mechanical twitch profiles from 51 MUs from 10 participants across 148 recordings were obtained.

#### Statistical Analysis

The data presented in this paper are intended to act as a proof of concept, showing what the set-up and processing pipeline can achieve. For each value and graph, the number of data points, *n*, is stated in the accompanying text. For all histograms and box plots *n* = 390.

## Results

From 12 participants we identified 51 MUs and, across all trials, we obtained 390 MUAMs and MU-STA curves. Figure 6 shows an example of the two signal modalities plotted together, as well as their STAs. For each MU activation, the velocity inside the MU motion domain changes in a synchronous manner. Figure 7 shows an example of the outputs for a trial with four identified MUs. The identified units have variable discharge rates (figure 7(a)), as is expected when the force level is close to the recruitment threshold of the MU. Figure 7(b) and figure 7(c) compare the shapes of the electrical STA and the MU-STA – the electrical STA is unique to a given MU while the MU-STAs are similar in this example. The MUAMs in figure 7(c) are truncated at approximately 10 mm as throughout the experiment no MUs were detected below this level, because of the relatively superficial detection volume of the surface EMG (Merletti and Muceli, 2019).

**Figure 6.**
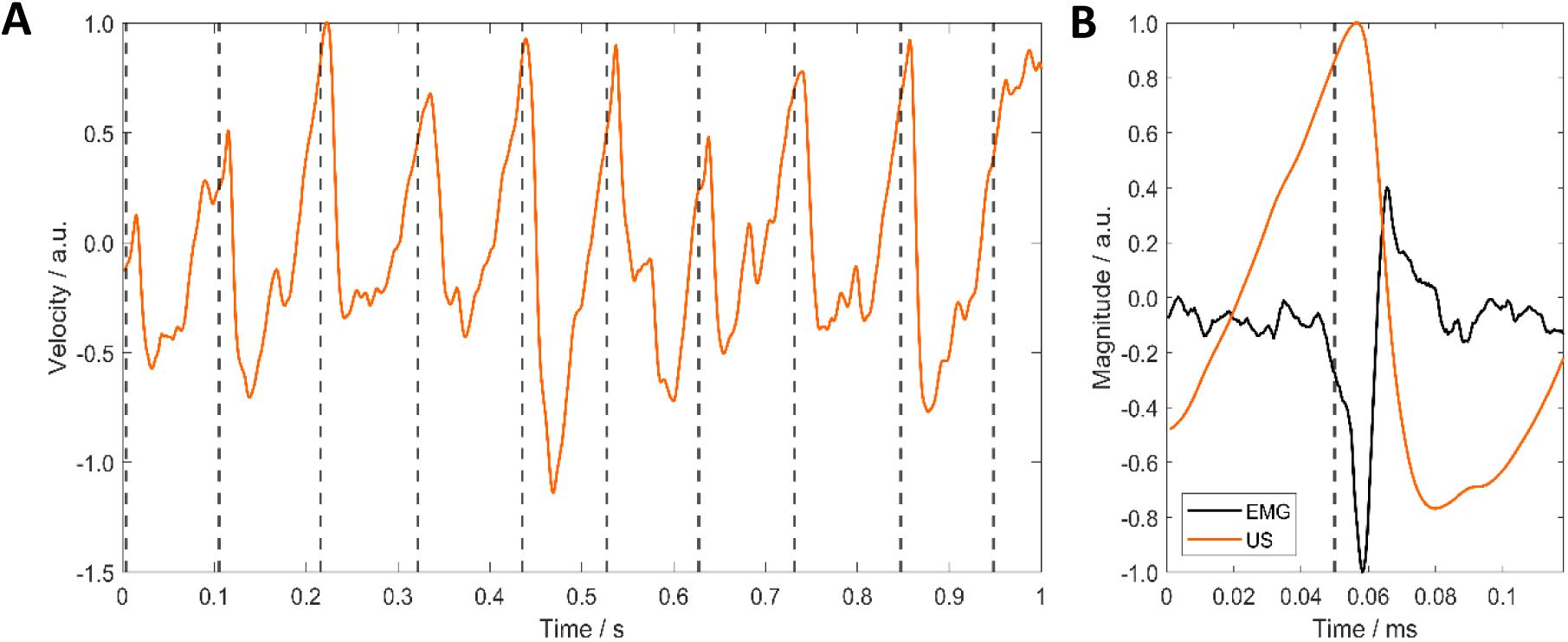
Synchronisation of velocity and EMG detected firing times. A) Example of average velocity inside motor unit (MU) motion domain for an identified MU (orange line), and action potential (AP) discharge times from EMG decomposition (black dotted lines) showing synchronicity. This is a 1 s interval extracted from a 30 s recording. B) Motor unit action potential (MUAP) found by means of spike-triggered average (STA) on the EMG signal (black line) and motor unit spike-triggered average (MU-STA) curve for the same MU (orange line). The black dotted line is the corrected firing time – the time at which the double differential of the highest signal to noise channel of the EMG array is above 5 standard deviations of the baseline noise level. The MU-STA is relaxing from the previous twitch hence the velocity is changing prior to the discharge time.

**Figure 7.**
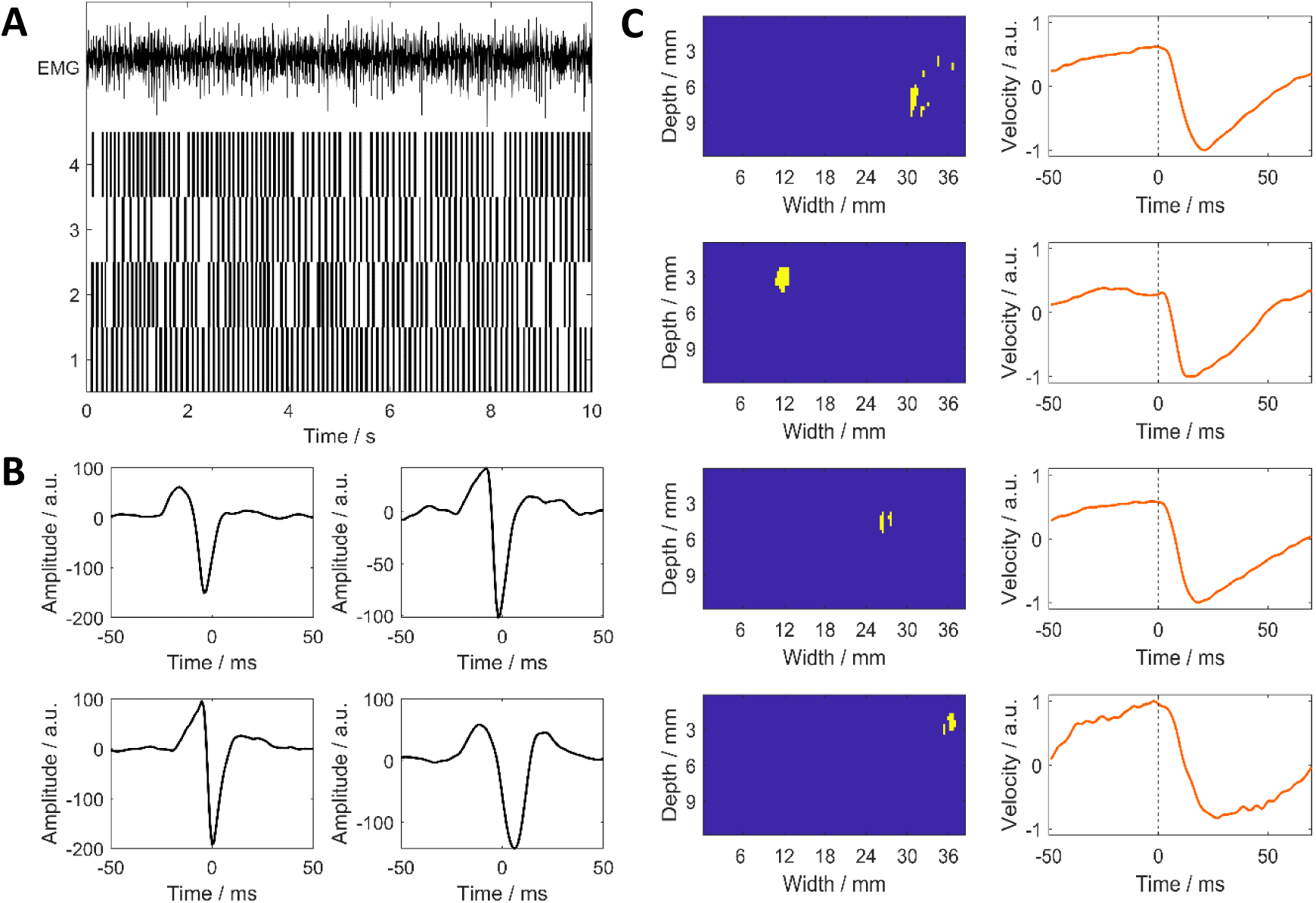
Example set of outcomes for a recording. A: Summed raw EMG signal (top) across the 64 recording electrodes in the high-density surface EMG array and the action potential (AP) discharge times of four of the identified motor units (MUs) – each of the units is given a label as shown to the left of the plot. A 10 s excerpt of the 30 s recording is shown. B: motor unit action potentials (MUAPs) for each of the identified MUs prior to correction to align activity onset, found by performing STA on the EMG signal. C: Motor unit activity maps (MUAMs) (left) and motor unit spike-triggered average (MU-STA) curves (right) for each of the four MUs. The MUAMs show the MU motion domains, and the MU-STAs show the average velocity in these regions in response to an AP. For all STA plots, the AP occurs at 0 ms.

### Motor Unit Spatial Motion Domain

To analyse the consistency of the area assigned as the MU motion domain across repeated trials, the percentage overlap of the spatial domains for each MU across different trials was calculated. The results are shown in figure 8 for each participant, and the average for all participants was 47.0 ± 17.4 %. In contrast, the percentage overlap between a given MU motion domain and the motion domains of other MUs was 6.9 ± 6.8 %. This showed that MU motion domains were more similar to the motion domains of the same MU in different trials than to those of other MUs for all participants. Although a 47% of overlap across trials may seem a low value, it incorporates error resulting from thresholding, as well as morphing and shifting due to other active MUs. The consistency in location of MU domain can be seen for representative MUs in figure 9.

**Figure 8.**
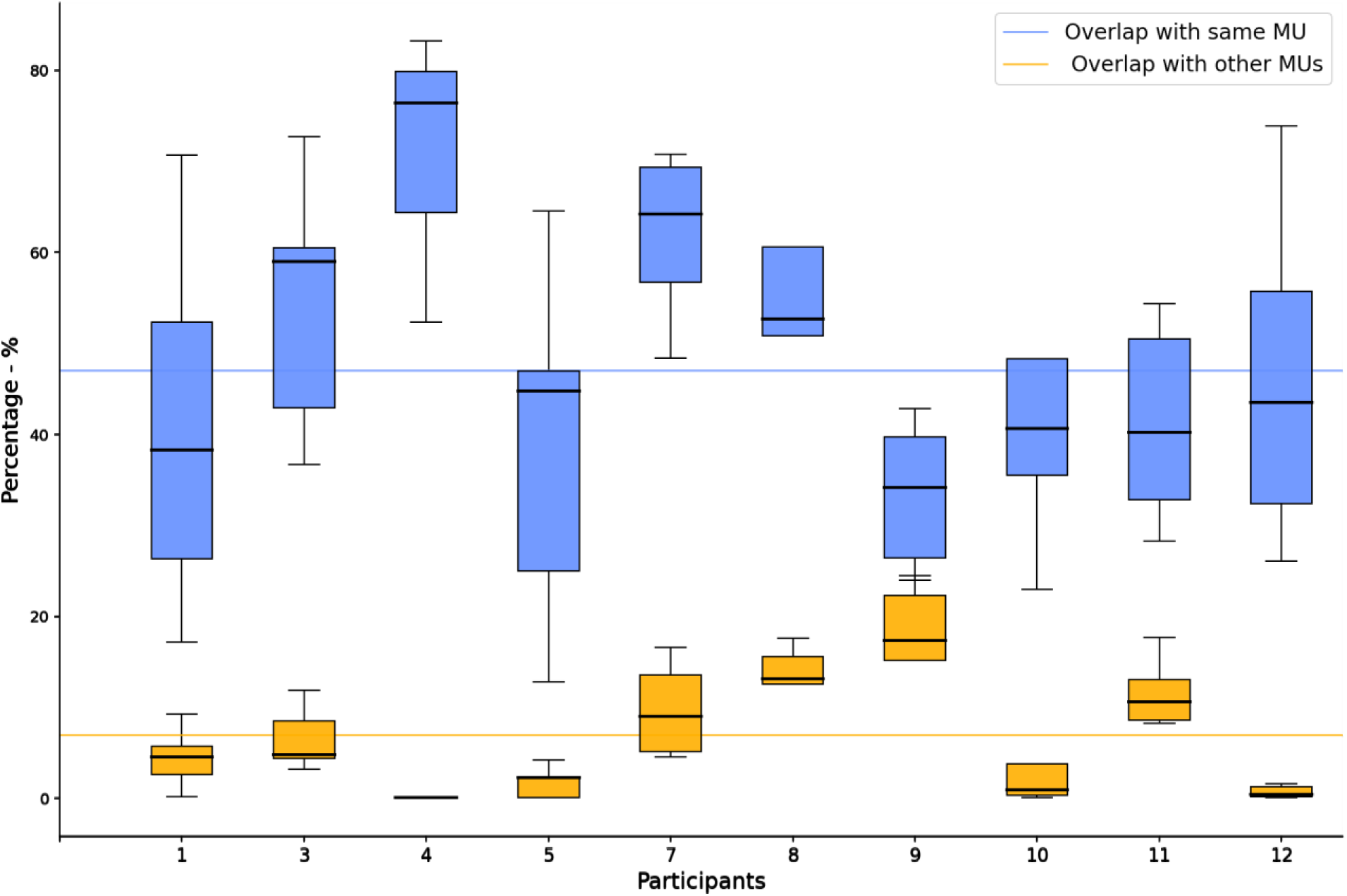
Box plot presenting participant-wise percentage overlaps of motor unit (MU) motion domains with other MU motion domains (orange), and with the same MU motion domains across different trials (blue). This is for 51 MUs identified in 390 trials across 10 subjects. For every participant the MU motion domains are more similar to the same MU motion domains in different trials than to other MU motion domains. This suggests that we are accurately identifying distinct MU motion domains from different simultaneously active units.

**Figure 9.**
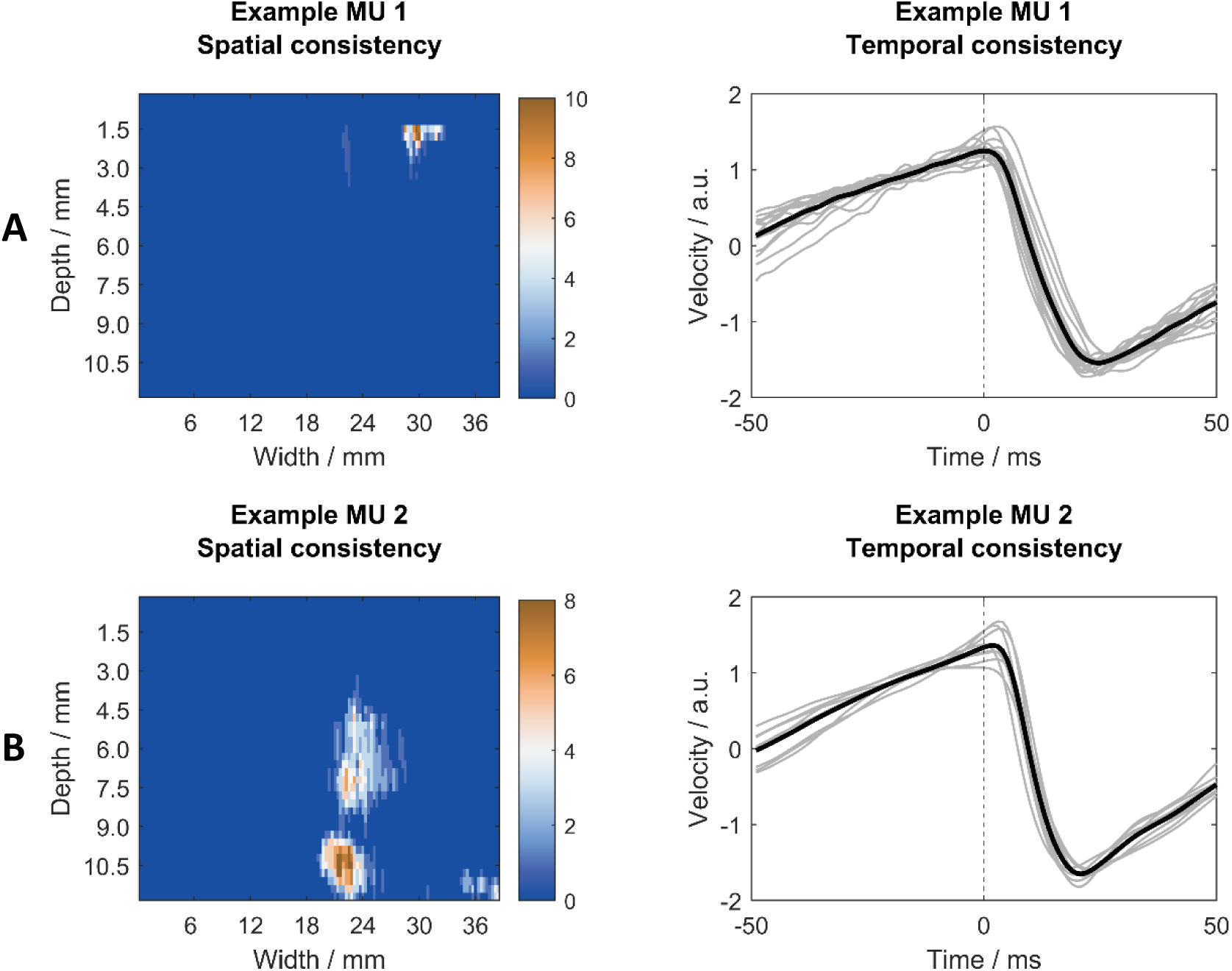
Repeatability across trials for MU identification. Left: motor unit activity maps (MUAMs) for two representative motor units (MUs) across 17 trials (panel A) and 8 trials (panel B) showing high consistency in location of MU motion domain, with some shift as expected due to differing activity level. Each plot is made by overlaying the MU motion domains for all trials containing that MU, such that pixels which are not in the motion domains for and trials have a value of zero. It can be seen from these plots that the regions are very consistent across trials, however there is shifting and morphing. Right: normalised motor unit spike-triggered average (MU-STA) curves for the same example MUs across all trials (grey lines) and average across these trials (black lines). These examples show the high repeatability in space and time.

### Mechanical Twitch Reproducibility

Figure 9 shows, for two representative MUs, both the MUAMs (left panel) and the normalised MU-STA curves (right panel) obtained for different trials (Example MU 1: 17 trials, Example MU 2: 8 trials). Each grey curve is the MU-STA velocity profile, averaged over the MU motion domain. These curves show high consistency across trials, despite changes in conditions (differing number of active MUs). For all 51 MUs across the 390 trials, the average standard deviations between curves from the same unit was 6.69 %. From these results, we concluded that the MU-STA curves were highly repeatable across force levels. Whilst it is clear that the MU motion domain morphs and shifts at differing force levels, the twitch profile is consistent across force levels, suggesting higher linearity in the temporal domain than in the spatial domain.

### Mechanical Twitch Duration, Contraction Time, and Activation Delay

Figure 10(a) shows the histogram of activation delays, with average value 3.8 ± 3.0 ms. A small proportion (3.8% of 390) of calculated activation delays were smaller than zero, suggesting an onset of mechanical motion prior to the electrical AP, due to numerical errors resulting from noise. Figure 10(b) shows a histogram of the mechanical twitch duration, with average value 17.9 ± 5.3 ms. Figures 10(c) and 10(d) show histograms for contraction times, with average values 12.1 ± 4.0 ms and 56.6 ± 8.4 ms for the active and total contraction times, respectively. All these measures are shown here for the first time for individual MUs in voluntary contractions.

**Figure 10.**
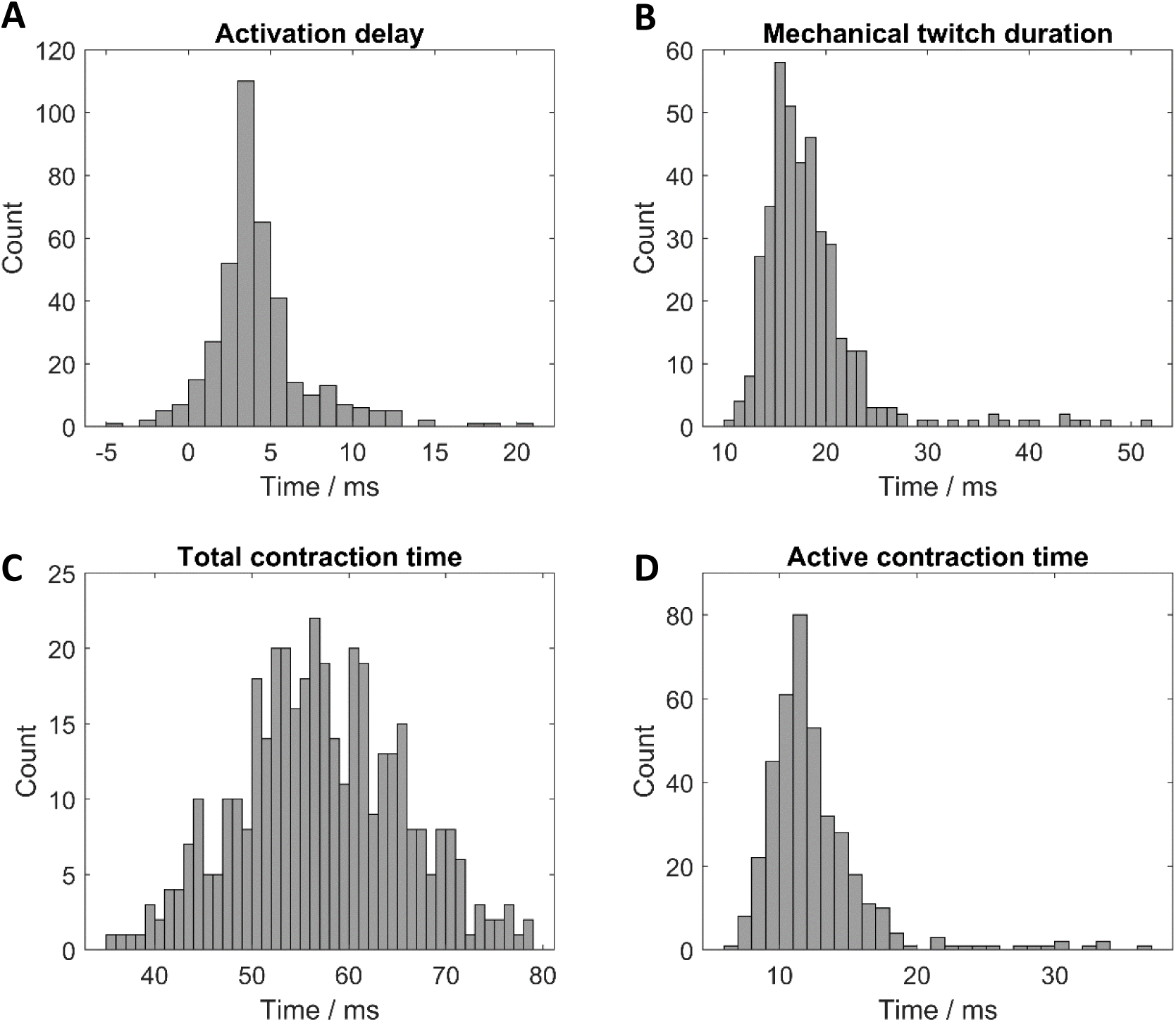
Histograms of temporal features extracted from the velocity profiles of motor unit twitches. A) Histogram of activation delay. B) Histogram of duration of mechanical twitch. C) Histogram of total contraction times. D) Histogram of active contraction times. For all histograms, *n* = 390 from 51 MUs across 10 participants.

### Muscle Twisting

In some instances, we observed regions of both positive and negative motion, aligned depth-wise, in response to AP discharges. On average, 17% of the MUs identified in the experiment across all participants exhibited this behaviour. These can be explained by a twisting of the muscle unit, where one group of muscle fibres moves towards the probe next to a group of fibres moving away from the probe. An example is shown in figure 11(b). This is the first observation of this phenomenon in natural contractions, and is in agreement with the twisting seen in electro-stimulated contractions (Deffieux *et al*., 2008). In the aforementioned study a fibre bundle with an unknown number of recruited MUs was activated by stimulation, thus twisting could not be reliably accredited to an individual MU. However here, single MUs are studied, and fibre bundle twisting is seen. As such, our results suggest that twisting can occur on an individual MU level. The source of this twisting is unclear and could be attributed to tendon attachment, to the surrounding connective tissue, or could be a fundamental contractile property of the MUs.

**Figure 11.**
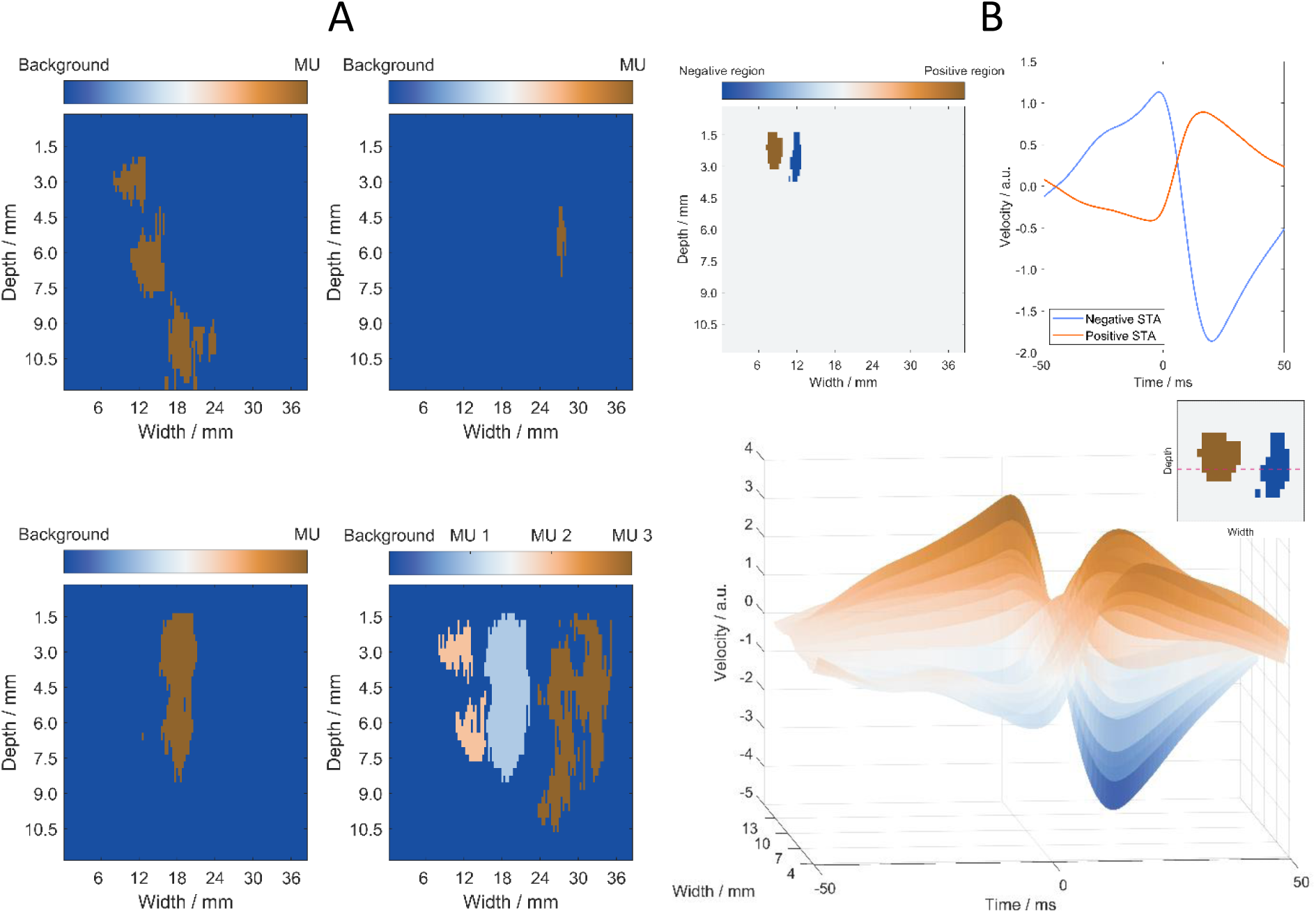
Examples of interesting motor unit (MU) behaviour. A) Different types of MU motion domain identified – these may be split, and vary hugely in size and shape. Also shown is a plot of the motion domains of 3 MUs active in the same trial (which don’t overlap in this case). B) An example of ‘twisting’ – part of the MU motion domain moves up and part moves down in response to a discharge. This is the first time this has been seen in natural neural contractions. Top: MUAM and MU-STAs for the positive and negative moving parts of the motion. Bottom: Mesh plot showing the motion along a line at a depth of 3 mm.

## Discussion

In this study we have proposed a method suitable for the kinematic study of individual MU twitch properties, with specific focus on extraction of E-C coupling time. This provides new perspectives in physiology for the study of muscular contractions at a granular level in relation to neuromuscular control. Furthermore, the approach could open up new avenues for the study and monitoring of pathologies that relate to muscle mechanics, and those which relate to the translation of motion from the electrical to the mechanical domains. Within this work, we have adapted the commonly used STA technique by adding a pixel-wise variability weighting to penalise inconsistency across spike occurrences, and thus minimise the contributions from pixels with large motion not associated with the spikes. This has implications for other applications where relevant signals may be masked, or where long recording times are not possible.

Current methods to study MU motion within a muscle suffer from several limitations. Often, these use electrical stimulation to elicit motion (Deffieux *et al*., 2008), evoking compound contractions rather than individual MU activity and resulting in contractions different to natural motor control. Further limitations include the use of systems with low temporal resolution compared to the durations of the involved physiological processes (Botter *et al*., 2013), low spatial resolution to properly separate contributions from multiple MUs (Birkbeck *et al*., 2020), or the deployment of invasive techniques, impacting the mechanics of the system and enabling the detection of just one MU at a time (Rohlén, Stålberg and Grönlund, 2020). The use of ultrafast US to decompose contributions to muscle motion from individual MUs has been previously attempted using spatio-temporal independent component analysis (Rohlén *et al*., 2020; Rohlén, Stålberg and Grönlund, 2020; Carbonaro *et al*., 2022), whilst we propose an approach which isolates these contributions by directly using the known MU discharge times.

### Technique validation

The online decomposition algorithm for the HDsEMG has been validated elsewhere, by comparison both with offline decomposition algorithms and with state-of-the-art iEMG techniques (Barsakcioglu *et al*., 2021). As such, by using these decomposition results to process the velocity maps, we ensure that our STA process identifies MU activity from a single unit. Furthermore, our STA processing involves dividing the velocity maps by the variance across each spike time, hence if no consistent movement was present the processed maps would be uniformly zero. The processing ensures that regions we are extracting are specific to that unit; the use of the division by the variance in the STA process is itself a validation that the extracted information is related to the single triggering units since otherwise the STA normalised by variance would tend to a null response.

Further, we validated the technique as a whole by analysing the repeatability of the MU-STA twitch profiles, and by comparing the MU motion domains across trials. The low standard deviations of the MU-STA curves found for the MUs across different trials showed the repeatability of the method, despite the *in vivo* nature of the work, and the variation in force level across different trials.

Spatially, MU motion domains were found to have higher similarity to the motion domains of the same MU in different trials than to that of different, simultaneously active, MUs (on average, 47% vs 7% respectively). The former shows high spatial consistency of MU motion domain location serving as evidence that we are correctly identifying consistent MU motion domains. As expected, this metric of consistency resulted in values lower than 100% due to factors such as probe and muscle shifting, translation and muscle morphing due to other active MUs, concomitant activation of different MUs, noise, and thresholding effects.

Although these values show that the method successfully identifies and separates contributions to muscular motion from multiple simultaneous MUs, they also highlight that even constant isometric contractions imply complex dynamics at the muscle unit level, resulting from mechanical coupling with neighbouring units. Further evidence for the validity of the technique is the agreement of some of the measured physiological parameters with the corresponding parameters identified with different approaches in previous studies, as discussed in the following section.

### Physiology of the muscle unit

It should be acknowledged that, due to the pennation angle of the muscle fibres in the TA, the cross-sectional images of the muscle do not correspond to the physiological cross-sectional area. As such, a bulging circular region of fibres is not expected to be seen. Nonetheless, the influence of this factor is presumably small because of the small pennation angle, as can be observed in figure 2. The actual region of motion is more complicated, due to the angle of the fibres as well as the compressibility of surrounding tissue and the effects of the pressure from the attached probe. However, this motion is still synchronised with the electrical events, and localised to a region.

Our analysis in the temporal motion domain yielded three mechanical twitch defining parameters: twitch duration, active contraction time, and total contraction time. In their work, Rohlén et al. identified a parameter called ‘twitch duration’ (approximately 50 ms) (Rohlén *et al*., 2020) and Deffieux et al. identified a parameter called ‘contraction time’ (70.1 ± 2.0 ms) (Deffieux *et al*., 2008), analogous to our ‘total contraction time’ (57.2 ± 8.7 ms). Our results are within one standard deviation of those by Rohlén et al. (Rohlén *et al*., 2020). Differences among studies may be due to the different muscles investigated, and methodological differences. On the other hand, larger differences in comparison to the results by Deffieux et al. (Deffieux *et al*., 2008) are likely due to their use of electrical stimulation.

From the time-domain analysis, we extracted the activation delay, i.e. the time between the electrical AP onset and the onset of motion caused by the twitch. We believe that the main contribution to this value is the E-C coupling, and this work therefore represents the first attempt to measure the duration of the E-C coupling in individual MUs in sustained voluntary contractions, which has potential impact for further study of neuropathies, myopathies and other conditions known to affect certain aspects of AP transmission (Orizio *et al*., 1997; Granata, Ikeda and Abel, 2000; Kaneko *et al*., 2002). In electrically stimulated contractions, the time between the stimulation and the onset of fascicle motion is attributed to the combination of E-C coupling and synaptic transmission (6.05 ± 0.64 ms, (Nordez *et al*., 2009)), 5.57 ± 1.37 ms (Hug *et al*., 2011)). Whilst these studies measure the time between electrical stimulation and fascicle motion, our study instead measures the time between onset of electrical activity as detected by HDsEMG and onset of fascicle motion, thereby not including the time for synaptic transmission. We would therefore expect smaller latencies in our study as it accounts for E-C coupling only. Another stimulation-based study instead measured the time between EMG onset and fascicle motion (2.2 ± 0.3 ms (Esposito, Limonta and Cè, 2011)), which therefore only accounts for E-C coupling, and is closer to our value of 3.8 ± 3.0 ms. It must be noted that the measurements of the activation delay require a very accurate synchronisation between electrical and mechanical measures as the delays are very small. For example, in previous work, the electrical activity seems to follow rather than precede the onset of mechanical activity (see figure 3 in (Rohlén, Stålberg and Grönlund, 2020)), which is likely due to issues in synchronisation or processing.

It is known that the EMD is more than two times greater for voluntary contractions than stimulated contractions, which is often attributed to the reversed recruitment order resulting in faster force development during stimulated contractions (Hopkins, Feland and Hunter, 2007). However, some studies also suggest that the E-C coupling is also greater in voluntary contractions than stimulated contractions (19.21 ± 6.79 ms (Begovic *et al*., 2014)), in contrast to our results which well align with those reported for stimulation studies. This may be explained by the use of bipolar EMG in these previous studies and by the influence of electrode location with respect to the innervation zones. For example, for electrodes placed at a distance of 4 cm from the innervation zone, there would be a detection delay due to propagation along the muscle fibres of approximately 10 ms (Hug, Lacourpaille and Nordez, 2011). Furthermore, the use of A-mode ultrasound to measure the muscle thickness and estimate the onset of fascicle motion may introduce errors due to muscle morphing and A-mode placement. This was circumvented in the current study by using B-mode ultrasound with a processing pipeline to identify individual MU motions. Our results suggest that E-C coupling is similar in stimulated and voluntary contractions.

Spatially, we defined a ‘MU motion domain’ as the region of muscle which moves with magnitude and synchronicity above a defined threshold when an AP travels along the muscle unit. Anatomically, MU territories have been defined as the cross-sectional area of the innervated fibres by a MU. MU territories have been studied using glycogen depletion experiments (Edström and Kugelberg, 1968; Garnett *et al*., 1979; Bodine *et al*., 1988) and scanning EMG (Stålberg and Dioszeghy, 1991). The latter method analyses the MU territory in one dimension, thus the measured size depends on the location of the needle intersection with respect to the MU cross-section (Gootzen, 1990). Furthermore, ‘silent zones’ (regions of no EMG signal) are seen in these measurements, which may be a result of the innervation being scattered in different portions of the muscle (Stålberg and Dioszeghy, 1991). More recently, MRI was paired with electrostimulation to measure human MU territories *in vivo* (Birkbeck *et al*., 2020). These studies found MU regions which they categorised as either circular, elliptical, crescent shape, spider, or split, corresponding well with our MU motion domain shape, which varied largely between MUs (figure 11(a)). The MRI experiments did not detect overlapping MUs due to methodology constraints, whereas we identified overlapping MU motion domains in every participant. On the other hand, split territories were not identified in a study using ultrafast US (Carbonaro *et al*., 2022), presumably because of methodological constraints. Our work therefore presents the first method able to detect both split and overlapping regions of activity.

### Limitations and Future Applications

While the work presented marks a novel approach to the study of MUs, there are some limitations of note. The results presented only show the technique working on low-force contractions (maximum 10% MVC). Translating the results to higher MVCs is not straightforward as they generate high global motion that obscures the individual MU discharges. Higher forces also increase muscle stiffness and increase the likelihood of probe movement and slippage. So far, the processing assumes that the probe remains stationary throughout the recording, which is a valid assumption for low forces; however, for higher forces (or dynamic contractions) this would not hold and considerations for its movement would have to be made.

Due to the use of HDsEMG to identify MUAPs, our methodology currently can only detect MUs near the surface of the skin. However, had we used iEMG we would detect a smaller number of individual units within close range of the needle tip (Lowery, Weir and Kuiken, 2006), and the needle (for the case of needle recordings) would cause mechanical coupling disrupting the results. In principle this can be solved by directly decomposing the US in MU discharges (Ali, Umander and Rohlén, 2020; Rohlén *et al*., 2020; Rohlén, Stålberg and Grönlund, 2020), however further validation on the basic assumptions underlying the mechanical components and coupling of MU twitches is necessary.

On the processing side, the autocorrelation velocity tracking algorithm employed is limited to one dimension, however the motion generated by the fibres contractions is multidimensional. Expanding this into a two-dimensional tracking so both forms of the fibre cross-sectional expansion movement can be tracked could be considered. On the same note, the resolution of our US image is limited so individual muscle fibres cannot be detected.

Finally, a formal method of selecting the threshold for producing the MUAMs could be implemented by, for example, optimising the threshold by comparison with the dimensions of MU as measured using scanning EMG. In another study, the spatial representation of the MUs in the biceps brachii appear to span a large portion of the muscle region (see figure 4 in (Carbonaro *et al*., 2022)), in contrast to our narrower domains. This discrepancy may be a result of the applied methodology in this previous study since EMG was used as a constraint for selecting the region, or of our use of an empirically chosen thresholding value.

## Conclusion

In conclusion, we have utilised synchronised HDsEMG with online MU decomposition and ultrafast US to isolate contributions to muscle motion from individual MU twitches. In doing so, we have been able to measure the E-C coupling time under non-stimulated conditions for individual MUs. This methodology and the results it yields open up avenues for monitoring pathology development and treatment, and could allow for further study of other features of the MU.

## Supplementary Information

Additional supporting information can be found in the online Supporting Information section. The available files are:

**Supplementary Video 1**. Spike-triggered average muscle motion due to an individual motor unit discharge.

## Declarations

## Acknowledgements

Figure 1 was created using BioRendr.com.

## Authors’ Contributions

All experiments were performed at the Department of Bioengineering at Imperial college London. E.L.: developed concept, wrote and developed code for control of Verasonics system, for synchronisation, and for post-processing and data analysis, designed custom probe holder, did in vivo experiments, analysed data, wrote paper. B.S.: developed concept, did in vivo experiments, analysed data, wrote paper. D.B.: developed concept, developed code for online EMG decomposition as well as closed-loop biofeedback framework and graphical user interface. D.F., M.T., J.I.P: developed concept. All authors revised the manuscript and provided intellectual feedback. All authors have read and approved the final version of this manuscript and agree to be accountable for all aspects of the work in ensuring that questions related to accuracy or integrity of any part of the work are appropriately investigated and resolved. All persons designated as authors qualify for authorship, and all those who qualify for authorship are listed.

## Availability of data and materials

As the datasets acquired are large, the data was not registered in a public database but may be made available, as appropriate and reasonable, upon request.

## Consent for publication

Not applicable.

## Ethics approval and consent to participate

All participants signed informed consent before participation in the study. Procedures and experiments were approved by the Imperial College Research Ethics Committee (ICREC reference: 20IC6422) in accordance with the declaration of Helsinki.

